# Unveil the Molecular Interplay between Aminoglycosides and Pseudouridine in IRES Translation

**DOI:** 10.1101/2024.09.20.614200

**Authors:** Yu Zhao, Chong Xu, Xin Chen, Hong Jin, Hong Li

## Abstract

Eukaryotic ribosomes are enriched with pseudouridine, particularly at the functional centers targeted by antibiotics. Here we investigated the roles of pseudouridine in aminoglycoside-mediated translation inhibition by comparing the structural and functional properties of the wild-type ribosomes and those lacking pseudouridine (*cbf5*-D95A). We showed that the *cbf5*-D95A ribosomes have decreased thermostability and high sensitivity to aminoglycosides. When presented with an internal ribosome entry site (IRES) RNA, elongation factor eEF2, GTP, sordarin, hygromycin B preferentially binds to the *cbf5*-D95A ribosomes during initiation by blocking eEF2 binding and stalls the ribosomes in a non-rotated conformation, further hindering translocation. Hygromycin B binds to the inter-subunit bridge B2a that is known to be sensitive to pseudouridine, revealing a functional link between pseudouridine and aminoglycoside inhibition. Our results suggest that pseudouridine enhances both thermostability and conformational fitness of the ribosomes, thereby influencing their susceptibility to aminoglycosides.

**Highlights:**

- Loss of pseudouridine increases cell sensitivity to aminoglycosides
- Pseudouridine enhances ribosome thermostability
- Hygromycin B competes with eEF2 for the non-rotated ribosome
- Hygromycin B deforms the codon-anticodon duplex

## Introduction

It has been well established that the widespread distribution of pseudouridine (λλ′), a C5-glycoside isomer of uridine, in ribosomal RNA (rRNA) contributes to ribosome synthesis, structures and translation (1–4). While maintaining identical base pairing interactions as uridine, pseudouridine provides the additional N1 imino proton for binding water (4–9) as well as for favorable base stacking (10–12). Loss of a subset or all pseudouridine in rRNA has been observed to cause widespread defects ranging from cell growth, ribosome maturation, to translation fidelity (2–5, 13). In human, mutations in the nucleolar pseudouridine synthase, dyskerin (*DKC1*), responsible for rRNA pseudouridylation are found in patients who suffer from the X-linked dyskeratosis congenita (X-DC) or its severe variant form, Hoyeraal-Hreidarsson (HH) syndrome (14). Translation of messenger RNAs containing the internal ribosome entry site (IRES) elements is especially impaired in cells containing the X-DC or HH-associated DKC1 mutants (15–17).

Earlier structural and biophysical studies indicate notable changes in ribosome structures and conformations upon pseudouridine loss, especially at the interfaces of domains and subunits (4, 5, 13). In hypopseudouridylated ribosomes bound with the hibernating factor Stm1, the head and the body move independently whereas in the wild-type ribosome, they move in a correlated fashion(5). When bound with the Taura syndrome virus (TSV) IRES and driven by the elongation factor eEF2 and GTP, The head of the hypopseudouridylated small ribosomal subunit exhibits swiveling motions exceeding those previously observed in yeast ribosomes (4). Furthermore, different from the wild-type ribosome, at the decoding center of the hypopseudouridylated ribosome, the A-site nucleotides, A1755 and A1756 of the small subunit helix 44 (h44) and A2256 of the large subunit helix 69 (H69) at intersubunit bridge B2a are stabilized in an unusual structure along with the bound eEF2 during IRES translocation (4). Strikingly, in human ribosomes, loss of only two pseudouridine (18S 609 and 863) led to increased efficiency in tRNA selection observed by single molecule Fluorescence Resonance Energy Transfer (smFRET) (13). These results consistently suggest that loss of the otherwise enriched pseudouridine isomers in ribosomes has a notable effect on the well conserved intersubunit bridge B2a that controls ribosome domain motions.

Recent studies reveal a fascinating link between chemical modifications in ribosomal RNA (rRNA) and the susceptibility of ribosomes to aminoglycosides, a potent class of antibiotics used against bacterial infections and some human diseases such as cancers and those associated with pre-termination (18). In both bacterial and eukaryotic ribosomes, aminoglycosides exert their activities primarily by binding to the decoding center of the ribosomes involving h44, H69 and the intersubunit bridge B2a where recognition of mRNA codons by transfer RNA (tRNA) or virus IRES is established (18–20). Single molecule FRET experiments show that binding of aminoglycosides can alter the conformational equilibrium of the ribosome in a chemical substituent-dependent manner (21–23). Notably, the decoding region is enriched with functionally important pseudouridine modifications, which aligns with the observed defects caused by their losses (4, 5, 13, 24). In yeast, removal of pseudouridine at three sites within the decoding region inhibited cell growth and elevated sensitivity to neomycin (25). Reduced pseudouridylation also resulted in an increase in sensitivity of mouse cells to sparsomycin and paromomycin while anisomycin rescued the growth defect (26). In E coli, elimination of the three conserved pseudouridine within H69 resulted in altered binding affinities of selected aminoglycosides for H69 as well as a different nucleotide protection pattern of H69 (24) . Whereas these results suggest an interplay between pseudouridylation and aminoglycoside activities, the molecular basis for this relationship remains lacking.

By using the previously established yeast strain (*cbf5*-D95A) that produces hypopseudouridylated ribosomes (5), we compared the susceptibility of the wild-type and the *cbf5*-D95A ribosomes to five aminoglycosides that target the decoding center. Cbf5 is the catalytic subunit of the box H/ACA small ribonucleoprotein particles (snoRNPs) responsible for installing more than 40 sites of pseudouridine. The Asp95 to alanine mutation of Cbf5 renders it inactive without impacting snoRNP assembly, therefore producing ribosomes that lack pseudouridylation (5). This strategy allowed us to assemble hypopseudouridylated ribosomes with TSV IRES during eEF2-mediated translocation in the presence of hygromycin B and analyze their cryo-electron microscopy (cryo-EM) structures. It further allowed us to study the thermal stability and the cell-free translation activities of the hypopseudouridylated ribosomes. Our structural and biochemical results provide insights into the binding mechanism of hygromycin B to the eukaryotic ribosomes during IRES initiation and how pseudourine regulates this process.

## Results

### Loss of Pseudouridine Increases Antibiotics Sensitivity

We employed spot assays to compare growth sensitivity to aminoglycosides targeting the decoding center between the wild-type and the *cbf5*-D95A cells. We first estimated the minimal inhibition concentration (MIC_50_) required to inhibit the growth of wild-type cells for each antibiotic by a combination of liquid and solid culture growth assays (Supplementary Figure 1a). A concentration around the MIC_50_ of each aminoglycoside was then applied in spot assays to compare sensitivities between the wild-type and the *cbf5*-D95A cells (Figure 1 & Supplementary Figure 1b). We also performed the spot assays at three different temperatures, 25 °C, 30 °C and 37 °C, respectively (Figure 1 & Supplementary Figure 1b). Finally, we included a non-ribosome targeting antibiotics, nystatin in this analysis to highlight the ribosome-specific effects (Figure 1 & Supplementary Figure 1b).

**Figure 1.**
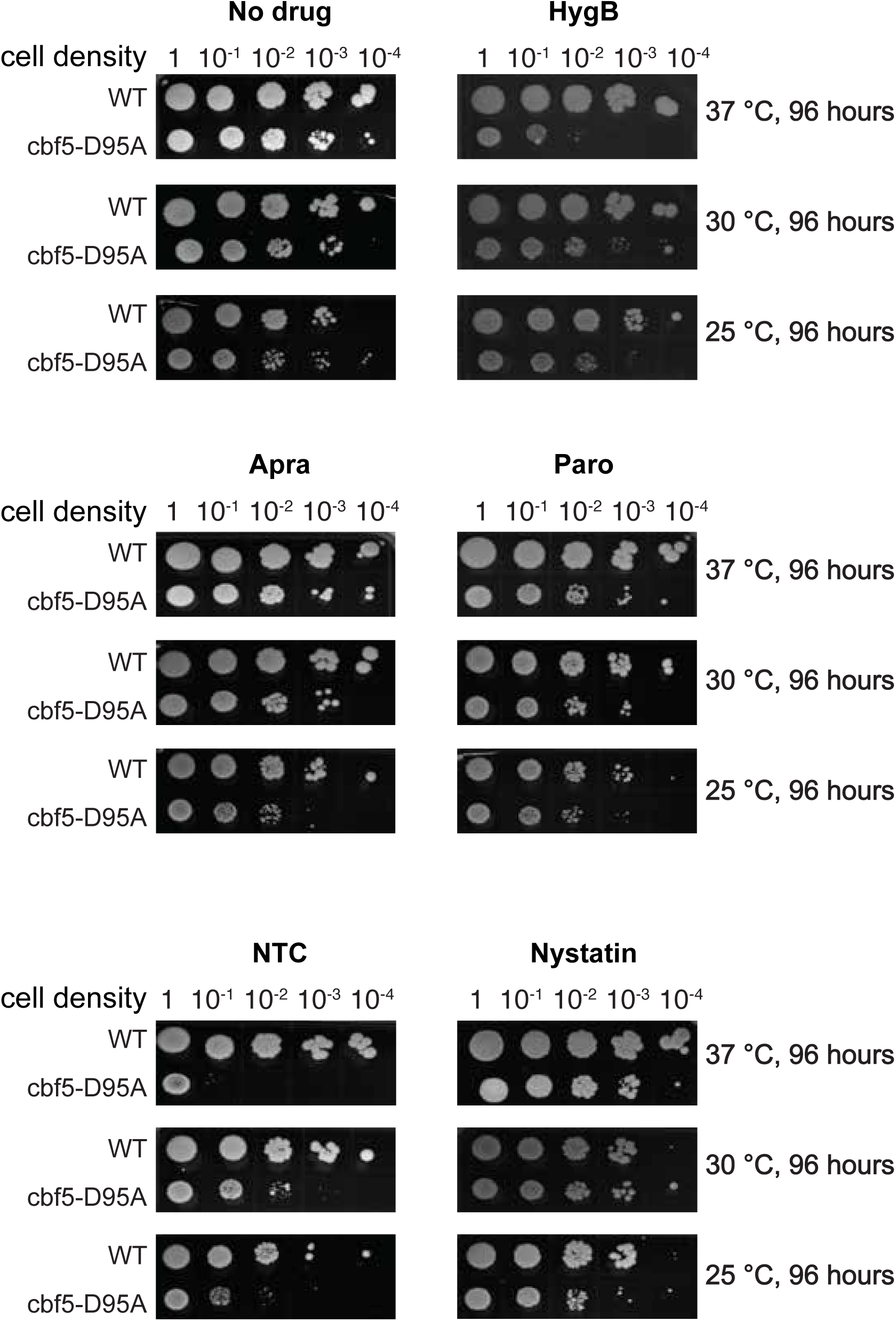
The *cbf5*-D95A cells are hypersensitive to translational inhibitors targeting the peptidyltransferase center (PTC). The growth on agar plates of yeast expressing either the wild type *CBF5* (WT) or *cbf5*-D95A mutant at three indicated temperatures over a period of 96 hours and different cell densities in the absence (No drug) or presence of translational inhibitors targeting the decoding center or nystatin (HygB, hygromycin-B at 15 mg/ml; Apra, apramycin at 0.5 mg/ml; Paro, paromomycin at 0.5 mg/ml; NTC, nourseothricin at 2 μg/mL; nystatin 10-33 μg/mL).

Unlike the wild-type, the *cbf5*-D95A cells exhibited significantly heightened sensitivity to the two broad-spectrum antibiotics, hygromycin B and nourseothricin (a natural mixture of streptothricins C, D, E and F), at all three growth temperatures (Figure 1a & Supplementary Figure 1b). Interestingly, the assay results further revealed an elevated sensitivity at 37 °C compared to 30 °C and 25 °C for the *cbf5*-D95A but not for the wild-type cells (Figure 1a & Supplementary Figure 1b), suggesting stronger inhibition of the hypopseudouridylated cells at elevated temperatures. Hygromycin B is a monophasic inhibitor of polypeptide synthesis (27) and binds to the decoding center of both bacterial and eukaryotic ribosomes (6, 28–30). By exerting an influence on the intersubunit bridge, hygromysin B efficiently impedes the translocation process of mRNA and tRNA (30–32). Less is known about nourseothricin mechanism of inhibition but in bacterial ribosomes, it targets multiple sites of the small subunit with a primary binding site at the interface between the head and the body domain (33). It is thus likely that nourseothricin targets similar sites of eukaryotic ribosomes.

The aminoglycosides examined are known to block translation elongation by targeting the universally conserved inter-subunit bridge B2a, which interferes with inter-subunit rotation dynamics (22) and tRNA translocation (23, 30, 34). The elevated sensitivity of the hypo-pseudouridylated cells to these inhibitors, therefore, suggests a higher potency of these antibiotics at inhibiting these processes, consistent with the altered translation fidelity and IRES-mediated translocation as a result of pseudouridine loss (4).

*Cbf5*-D95A cells also display enhanced susceptibility to other bacterial aminoglycosides, paromomycin and apramycin. However, unlike the pronounced temperature-dependent sensitivity observed for hygromycin B and nourseothricin (Figure 1), these antibiotics do not exhibit a similar effect, emphasizing the distinct inhibition mechanisms of different aminoglycoside classes and the role of pseudouridine.

### Pseudouridine Enhances Ribosome Thermostability

The high sensitivity of the *cbf5*-D95A ribosomes to aminoglycosides at elevated temperatures suggests a possible change in their thermostability upon pseuodouridine loss. We subsequently performed circular dichroism (CD) spectroscopy with both the *cbf5*-D95A and the wild-type ribosomes. We collected CD scans from 230 nm to 320 nm of the purified ribosomes as they were denatured by the increasing temperature. We used the ellipticity values at 260 nm as a measure of the RNA tertiary structure and plotted the RNA melting curves to obtain the melting temperature T_m_ (Figure 2). The melting curves were also plotted for ellipticity values at 230 nm for comparison as this region is mostly contributed by proteins (Supplementary Figure 2a).

**Figure 2.**
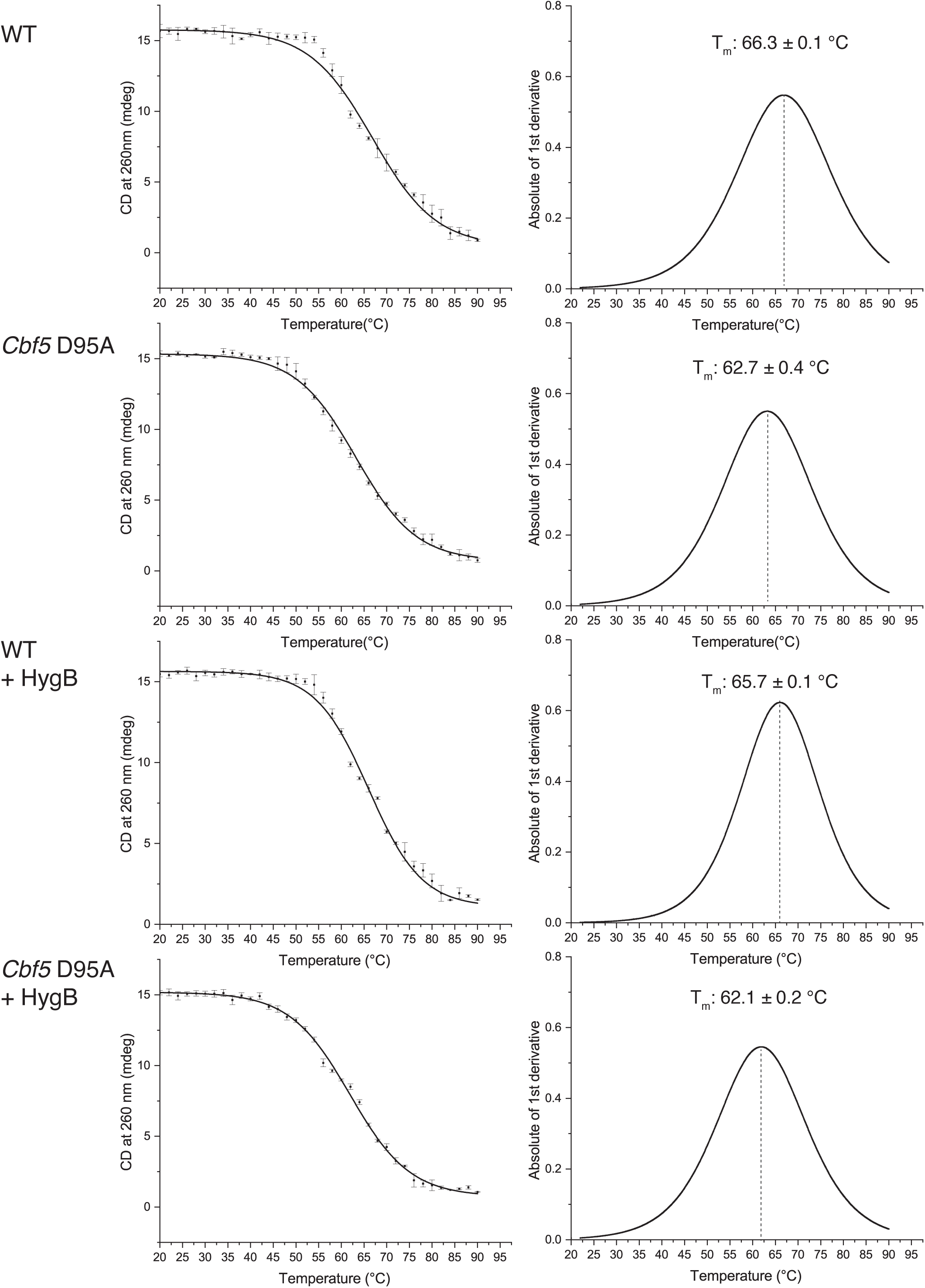
The *cbf5*-D95A ribosomes have decreased thermostability. The melting curves extracted from the ellipticity at 260 nm of the circular dichroism (CD) scans (left) and the first order derivative of the melting curves (right) for the wild-type and the *cbf5*-D95A ribosomes in 100 mM KCl, 25 mM HEPES pH7.5, 2 mM DTT, and 10mM MgCl_2_. The calculated the melting temperatures are indicated as mean ± standard deviations. Three experimental replicas were performed from which the standard errors were computed.

Whereas the wild-type ribosomes began denaturing at 53°C, the *cbf5*-D95A ribosomes started at a lower temperature, 42°C, leading to their calculated melting temperatures of 66.3°C+/-0.1°C and 62.7°C+/-0.4°C, respectively (Figure 2a). Interestingly, experiments with isolated RNA molecules such as the H69 model showed more drastic difference in T_m_ between the modified and the unmodified RNA (10), suggesting that the bound proteins and the other molecules of the intact ribosome can partially compensate the difference. The T_m_ measured from ellipticity at 230 nm showed no noticeable changes between the two ribosomes, suggesting the stability decrease is largely attributable to RNA. Consistently, reduced magnesium lowered the T_m_ of both the wild-type and *cbf5*-D95A ribosomes but to a similar degree (Supplementary Figure 2b). Furthermore, while isolated RNA fragments can be substantially stabilized by antibiotics binding (35), molar excess hygromycin B did not change the melting temperature of the wild-type (65.7 +/-0.1 °C) or the *cbf5*-D95A (62.1+/-0.2°C) ribosomes (Figure 2a).

The consistently lower T_m_ of the *cbf5*-D95A than that of the wild-type ribosomes under various conditions revealed reduced stability of the ribosomes assembled with the rRNA without pseudouridine. The direct impact of pseudouridylation on ribosome thermostability reflects the previously observed changes in solvent structures and domain motions in ribosomes upon pseudouridine loss (4, 5).

### Hygromycin B Stabilizes Non-rotated Decoding Structure

In our previous study of the *cbf5*-D95A ribosomes during IRES-mediated initiation in absence of hygromycin B (4), we observed rearranged decoding center, especially in the otherwise heavily pseudouridylated H69. Given that hygromycin B binds to the decoding center, we obtained cryoEM structures of the ribosomes isolated from the *cbf5*-D95A cells assembled with TSV IRES, eEF2, GTP, sordarin, and hygromycin B (Supplementary Figure 3 & Supplementary Table 1).

After extensive 3D classification, particles are sorted into two major classes, one without eEF2 (initiation) and one with eEF2 (translocation) (Supplementary Figure 3, Supplementary Figure 4 & Supplementary Table 1). It is immediately clear that hygromycin B binds only to the initiation complex (or back translocated upon eEF2 departure) and not to the translocation complex (Figure 3), suggesting that hygromycin B primarily interferes with the processes prior to eEF2 binding. The presence of eEF2 thus overcomes the effect of hygromycin B inhibition, consistent with the mechanism described biochemically (31). Since the hygromycin B-free structures have been previously characterized (4), we focused on the hygromycin B-bound initiation complexes (Figure 3a, Supplementary Figure 3, Supplementary Figure 4 & Supplementary Table 1).

**Figure 3.**
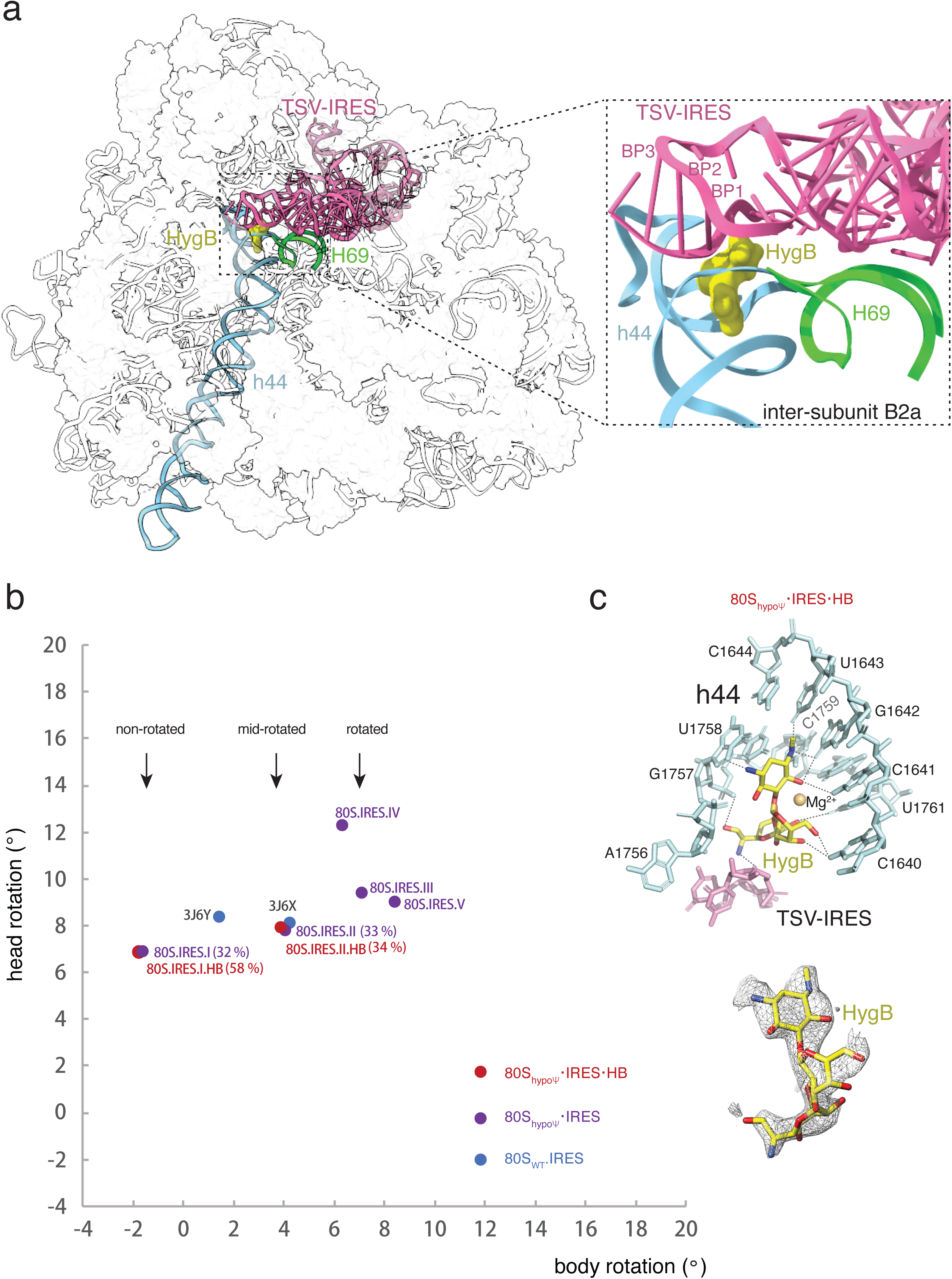
Structures of hygromycin B bound to the TSV-IRES initiation complex. **a.** CryoEM structure of *cbf5*-D95A ribosomes bound with TSV-IRES (pink) and hygromycin B (HygB, yellow). The helix 69 of the large subunit (H69) is colored in green and the helix 44 of the small subunit (h44) is colored blue. Inset shows a close-up view of the bound hygromysin B wedged into a pocket formed by TSV-IRES, h44 and H69**. b.** The subunit rotation map of the TSV-IRES initiation complex of the wild-type ribosome (80S_WT_•IRES), the hypopseudouridylated ribosome without hygromycin B (80S_hypoψ′_•IRES) and the hypopseudouridylated ribosome bound with hygromycin B (80S_hypoψ′_•IRES•HygB). Each filled circle represents a set of head (with respect to body) and body rotation (with respect to the large subunit) angles. The precent of particles contributed to each conformation is indicated in parenthesis. The wild-type ribosome conformations are from the structures indicated by their PDB IDs. The approximate positions of the non-rotated, mid-rotated and rotated head are marked by arrows. **c.** The interactions (top) and density of the bound hygromycin B. The h44 nucleotides are shown in light blue and are marked. The IRES nucleotides are shown in pink. The bound hygromycin B is colored in yellow. Dashed lines indicate close contacts between hygromycin B and the RNA nucleotides.

The hygromycin B-bound initiation complexes largely populate two states, the non-rotated (structure I, 80S_hypoψ′_•IRES•HB.I, ∼58%) and the rotated (structure II, 80S_hypoψ′_•IRES•HB.II, ∼32%) state (Figure 3b, Supplementary Figure 3 & Supplementary Table 1). Considering the body rotations, structure I, in its non-rotated state, resembles the classical state of the ribosome bound with mRNA and tRNA. In contrast, structure II is similar to the hybrid state. The dynamic exchange between the two conformations is essential for translocation, where the hybrid state is believed to serve as the substrate for elongation factor binding (36). Both the TSV IRES-bound wild-type (37) and hypopseudouridylated(4) ribosomes also populate the two similar states that are, however, equally populated (Figure 3b). In contrast, the hygromycin B-bound initiation complex prefers the non-rotated state (Figure 3b). The preferential population of the non-rotated state depletes the hybrid state, effectively halting eEF2-mediated translocation. This mechanism of inhibition is similar to that established for bacterial ribosomes in which hygromycin B inhibits the spontaneous reverse translocation (30). The decreased RNA stability due to pseudouridine loss likely amplifies the effect, making the ribosome more susceptible to hygromycin B inhibition.

### Hygromycin B Occupies Diphthamide Binding Site and Deforms Codon Nucleotides

In both initiation complexes, structures I & II, hygromycin B binds at the intersubunit bridge B2a consisting of h44 of 18S and H69 of 25S (Figure 3c & Supplementary Figure 4), similar to the binding mode observed in the hygromycin B-bound wild-type human ribosome (6) with one exception. Unlike the human h44-equivalent nucleotide of A1755 (A1824) that stays in helical stack and forms close contacts with hygromycin B (6), A1755 flips out and forms no contact with hygromycin B (Figure 3c & Supplementary Figure 4). The bound hygromycin B otherwise forms an extensive network of interactions with other h44 nucleotides 1757-1759 and 1640-1643 as well as IRES nucleotides 6949-6951 mimicking the A codon (Figure 3c & Supplementary Figure 4). The direct contact between hygromycin B and the codon-like nucleotides is unique among aminoglycosides. Other aminoglycosides primarily engage only with h44 or other rRNA nucleotides (22). We did not observe any secondary hygromycin B binding site as seen in bacterial ribosomes (29). We also did not observe the base stacking between the yeast-equivalent bacterial A site nucleotides A1492 of h44 and A1913 of H69 when hygromycin B is bound to bacterial ribosomes (30), suggesting a difference between the prokaryotic and the eukaryotic ribosomes.

Superimposition of the translocation and the initiation complex structures showed that the bound eEF2 in the translocation complex would clash with the bound hygromycin B in the initiation complex (Figure 4a). Specifically, the ring 4 moiety of hygromycin B occupies the same site of the decoding center as the diphthamide residue His699 of eEF2 (Figure 4a). Diphthamide is critical in stabilizing the codon-anticodon duplex in both cap- and IRES-dependent translation (38, 39) and its structural environment is sensitive to pseudouridine modification (4). Similarly, the ring I motif of hygromycin B establishes close contacts with the IRES nucleotides mimicking codons (Figure 3c). The overlap between hygromycin B and diphthamide signifies a similar structural strategy used by both to influence translation, although with contrasting consequences.

**Figure 4.**
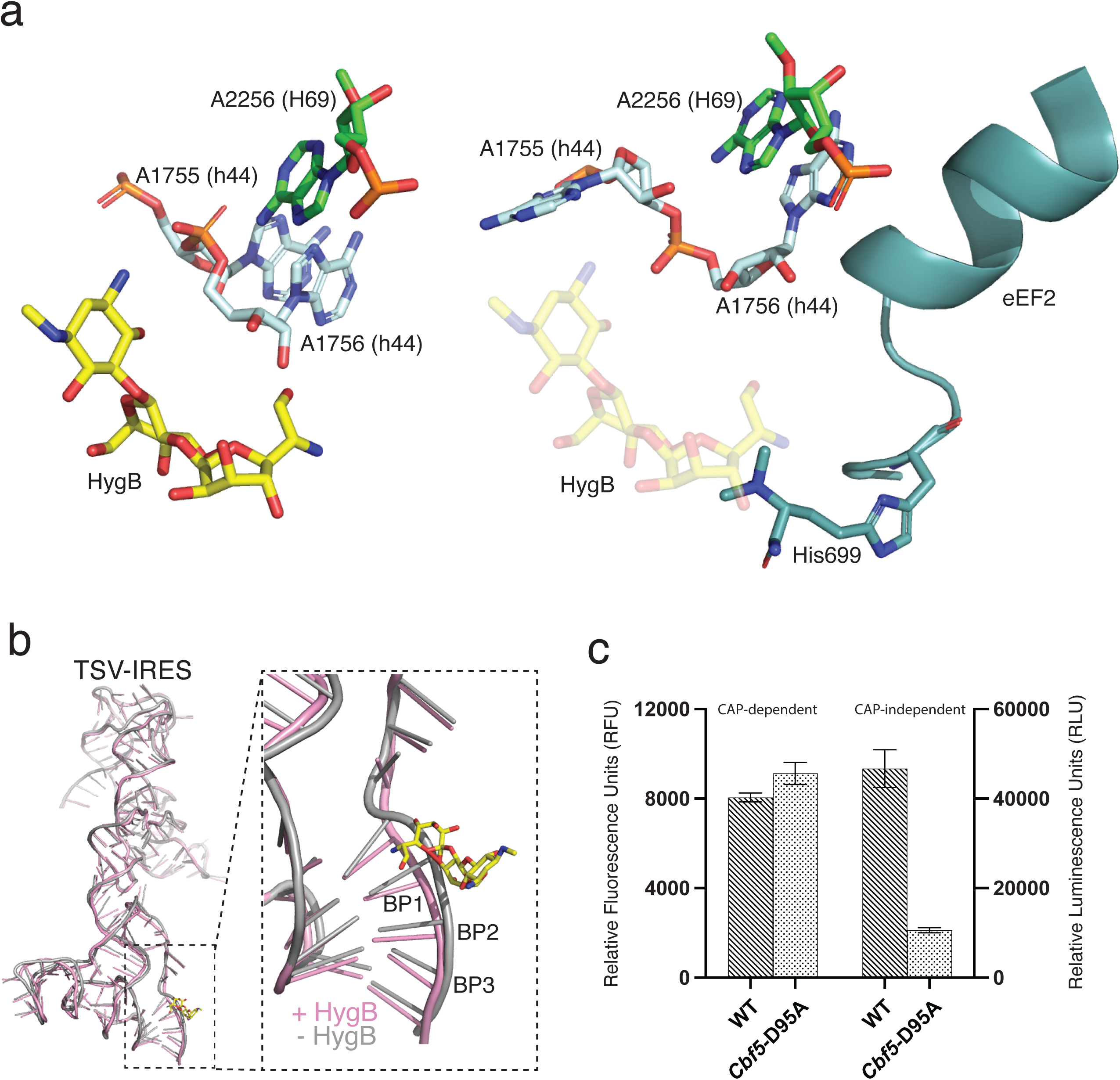
Inhibition mechanism of hygromycin B on IRES translocation and the role of pseudouridylation. **a**. Hygromycin B (HygB) competes with eEF2 binding. Comparison of the TSV-IRES initiation complex structure (left) and the translocated structure (right, PDB 8EUB) reveals the clash between the bound hygromycin B (yellow) and the diphthamide 699 of eEF2 (teal). **b.** Hygromycin B restructures the messenger RNA. Comparison of the initiation complex in the presence (pink) and the absence (gray) of hygromycin B (yellow) reveals a clash between hygromycin B and the first base pair of the codon (BP1) at the A site. **c**. The reporter assay is used to detect changes in the cell-free translation activities caused by the loss of pseudouridine of the *cbf5*-D95A ribosome. Relative fluorescence levels, indicating cap-dependent translation, were measured using eGFP signal. Relative luciferase activity, corresponding to cap-independent translation, was measured using NanoLuc with the Nano-Glo Luciferase Assay System (Promega). Both fluorescence and luciferase activities were measured after the cell-free in vitro translation reaction. Data are presented as the mean ± SD of three independent experiments.

Comparison of the TSV IRES structures in the initiation state with (structure I or structure II) and without (4) hygromycin B reveals a deformed codon-anticodon duplex by the aminoglycoside, regardless the domain rotational status (Figure 4b). Hygromycin B binds directly to the IRES nucleotides 6949-6952, displacing the codon nucleotides downward to create a binding pocket for itself. Despite minimal disruption to the remaining IRES structure, this localized structural change likely weakens IRES binding and the observed changes in ribosome domain conformations. Importantly, loss of pseudouridine amplifies these structural changes, further obstructing IRES binding and translation.

The influence of HgyB on *cbf5*-D95A ribosome structure is consistent with its vulnerability to the loss of pseudouridylation (4, 5). In both yeast and human cells, IRES-mediated translation depends on the presence of pseudouridylation activity (15, 26). To determine whether the compromised translation activity stems specifically from the *cbf5*-D95A ribosomes, we conducted cell-free translation assays with purified ribosomes from both wild-type and *cbf5*-D95A cells (Figure 4c). Notably, *cbf5*-D95A ribosomes exhibited normal activity in CAP-dependent translation, but a significant reduction in CAP-independent translation compared to wild-type ribosomes (Figure 4c). This finding suggests that pseudouridylation is especially critical for IRES-mediated translation, a process that relies more heavily on ribosome dynamics than CAP-dependent translation.

## Discussion

Though overwhelming evidence support an essential role of pseudouridine in ribosome structure and function, how this modification impacts ribosome thermostability as well as aminoglycoside activities remains unexplored. We analyzed the thermostability of the purified pseudouridine-free and the wild-type ribosomes by circular dichroism, which revealed a critical role of pseudouridine in maintaining the thermostability of the ribosome. We performed systematic analysis of cell growth in presence of both ribosome- and non-ribosome-targeting antibiotics that revealed increased sensitivity to aminoglycosides when ribosomes lack the pseudouridine modification. We subsequently obtained high-resolution cryoEM structures of hypopseudouridylated ribosomes bound with the aminoglycoside hygromycin B during TSV-IRES initiation and unveiled hyperstabilization of the ribosome at a state that hinders translocation. Our results consistently show that pseudouridine increases stability and fine-tunes the conformational dynamics of ribosomes.

The thermodynamic and structural roles of pseudouridine have previously been observed in model RNA (10–12). The observed impacts on the intact ribosomes are consistent with and further advance the importance of the pseudouridine modification. Importantly, our results highlight the unexpected allosteric effects of pseudouridine on the movement of ribosome domains, providing the structural and thermodynamic basis for the increased sensitivity to antibiotics.

The high sensitivity to the group of aminoglycosides targeting the decoding center coincides with it being both the site of conformational change and with highly enriched pseudouridine. Notably, H69 of the large subunit contains four well-conserved pseudouridines (Ψ2258, Ψ2260, Ψ2264, and Ψ2266) and supports the extrahelical adenosine A2256 important to decoding. We showed previously that loss of pseudouridine causes changes in H69 nucleotide conformations and hyperstabilizes its interaction with the elongation factor eEF2 during TSV IRES translocation (4), consistent with its impact on antibiotic binding at the same site.

The observation that mutations in ribosome biogenesis factors unrelated to chemical modifications can also lead to increased antibiotic sensitivity (40) suggests that this effect might be partly due to changes in the overall abundance of ribosomes, rather than solely attributable to alterations in chemical modifications. Despite so, the structural and biochemical evidence presented highlight the effects of pseudouridylation on the ribosome itself.

## Materials and Methods

### Cloning, Expression, and Purification of Yeast eEF2

Cloning and purification yeast eEF2 was similar as previously described(4). Briefly, eEF2 fused with a C-terminus 6x-his tag was purified from yeast cells transformed with pRS415-Gal-eEF2-6xHis vector, grown to OD600=0.6∼0.8 and induced by the addition of galactose. Nickel affinity was first used followed by MonoQ ion exchange and gel filtration chromatography. The purified eEF2-6xHis was concentrated, flash frozen and stored in buffer (25mM HEPES-KOH, 100mM KCl, 2mM MgCl_2_, 2mM DTT, pH 7.5) in aliquots at -80 °C.

### Spot Assay

For spot assays, Sc strain BY4741 cells harboring the *cbf5*-D95A point mutation were grown in YPD in presence of ampicillin (100 μg/mL) to saturation. After washing three times with cold sterile water, cells were diluted to 1 optical density (OD_600nm_)/mL, from which four 1-10 serial dilutions were made. The diluted cells were spotted onto agar plates containing appropriate concentrations of antibiotics (hygromycin B at 15 μg/mL, apramycin at 0.5 mg/mL, paromomycin at 0.5 mg/mL, nourseothricin at 2 μg/mL, and nystatin 10-33 μg/mL). The non-ribosome targeting antibiotics nystatin was used as a control. The agar plates were incubated for a total of 96 hours at 25 °C, 30 °C, and 37 °C, respectively, and were monitored at 36, 72 and 96 hours.

### Liquid Culture Assay

To determine MIC_50_ (Minimum Inhibitory Concentration at which 50% of growth is inhibited) for each aminoglycoside, a single colony of the wild type or the mutant cells was used to start growth in YPD containing varying concentrations of the antibiotics for 24 to 48 hours at 25 °C. The growth of the microbial cultures was monitored continuously by optical density measurement at 600 nm. The MIC_50_ was estimated by the lowest concentration of the antibiotics that inhibited the growth by 50% of that in the absence of the antibiotics.

### Melting Temperature Measurement

Melting temperature measurement was conducted by following circular dichroism (CD) spectrum changes upon thermal denaturation(41) using a Chirascan V100 Circular Dichroism spectrometer with a 1 mm pathlength quartz cuvette. Purified wild-type and *cbf5*-D95A ribosomes were diluted to 11 optical density (OD_260nm_)/ml in a CD buffer (100 mM KCl, 25 mM HEPES-KOH, 2 mM DTT, and 10 mM MgCl_2_) and cooled on ice for 20 minutes. Denaturation curves were generated by scanning ellipticity from 230-300 nm as temperature increased from 20 °C to 90 °C at a rate of 2 °C/min. The ellipticity at 260 nm was extracted for each temperature and plotted as a function of the temperature. The melting temperature was obtained by taking the first derivative of the melting curve (Origin Pro 2022). Three experimental replicas were obtained for each sample and errors were calculated as standard deviation.

### Purification of cbf5-D95A ribosome subunits

Ribosomes from the *Saccharomyces cerevisiae* strain BY4741 harboring the *cbf5*-D95A point mutation were purified similarly as previously described (5, 42, 43). Briefly, ribosomes obtained under the glucose starvation condition were incubated in splitting buffer (20 mM HEPES-KOH, 500 mM KCl, 5 mM MgCl_2_, 1 mM puromycin, 2 mM DTT, pH7.5) supplemented with RNase inhibitor for 1 hour at 4 °C and loaded on a 10%-30% sucrose gradient for extracting subunits by centrifugation. The subunits were pooled and buffer exchanged to a grid-making buffer (25 mM HEPES-KOH, 100 mM KCl, 10mM MgCl_2_, 2 mM DTT, pH7.5) and flash frozen and for storage at -80 °C.

### In vitro transcription and purification of TSV-IRES

The TSV-IRES mRNA 6741-6990 was obtained by T7 in-vitro transcription from a PCR product amplified from the plasmid containing the mRNA clone and the transcript was purified by using RNA purification kit (NEB) (4).

### Cell-free translation assay

Yeast cells were harvested at an OD600 value of approximately 0.6-0.8. The cell pellet was resuspended and lysed in Buffer A (30 mM HEPES pH 7.4, 100 mM KOAc, 2 mM Mg(OAc)_2_, 2 mM DTT, 466 mM mannitol and Roche Complete Protease Inhibitor Cocktail tablet). Cells were lysed in liquid N_2_ and the lysate was exchanged into Buffer B (30 mM HEPES pH 7.4, 100 mM KOAc, 2 mM Mg(OAc)_2_, 2 mM DTT, 20% glycerol and Roche Complete Protease Inhibitor Cocktail tablet) using a PD-10 G25 column. The absorbance at 260 nm (A260) was measured for each 0.5 mL elution fraction, retaining only those with an A260 exceeding 100. Subsequently, ribosomes were pelleted by ultracentrifugation in 4.0 mL polycarbonate thick-wall tubes at 100,000 g for 4 hours at 4°C using a 45Ti rotor. The supernatant from wild-type cells was collected as S100. Both wild-type and mutant cell ribosome pellets were resuspended in Buffer C (30 mM HEPES pH 7.4, 100 mM KOAc, 2 mM Mg(OAc)_2_, 2 mM DTT) and all samples ribosome solution were diluted to the same concentrations measured by A260 (∼150). Reporter DNA (Addgene #127332) was transcribed in vitro using the HiScribe T7 Kit with CleanCap (NEB), replacing partial GTP with m7G(5’)ppp(5’)A RNA cap structure analogous to the 5’ capped mRNA. The prepared mRNA was refolded by heating at 65°C for 3 minutes, then cooling on ice for

30 minutes. The mRNA was then mixed with S100 and ribosomes at equal RNA concentrations. The reactions were performed in buffer D (20 mM HEPES pH 7.6, 120 mM KOAc, 2 mM Mg(OAc)_2_, 1 mM ATP, 0.1 mM GTP, 20 mM creatine phosphate (Invitrogen), 0.01 mM of each of the twenty amino acids, 2 mM DTT, 0.12 U/μL creatine kinase (Invitrogen), 1 U/μL RNase inhibitor (NEB)). The reactions were incubated at 24°C for 2 hours in the PCR tubes. Luciferase activity was subsequently measured using the Nano-Glo Luciferase Assay System (Promega) and fluorescence activity was measured directly, both using a Synergy HTX plate reader. The data were visualized and processed using the R package ggplot2. P-values for significant levels were calculated using a t-test in R.

### Assembly of Inhibition Complex of 80S-IRES with hygromycin B

The 40S subunit at 200 nM concentration was incubated with 2 μM refolded TSV-IRES RNA for 10 min at 30 °C, followed by the addition of 200 nM of 60S subunit and another 10 min incubation, to form the 80S-TSV-IRES initiation complex. A premixed solution containing 4 μM of Hygromycin B, 1.7 μM of eEF2, 0.3 μM of sordarin, and 0.3 μM of GTP was added to the 80S-TSV-IRES initiation complex followed by additional 10 min incubation at the same temperature. The inhibition reaction mixture was then put on ice before making cryoEM grids.

### Cryo-EM Grid Preparation, Data Collection and Analysis

4 μl of the inhibition reaction mixture was applied onto plasma cleaned holey carbon grids coated with 2nm thick amorphous carbon (Quantifoil R 2/2, UT, 300 Mesh, Copper). Following 30 sec of incubation inside the chamber of FEI Vitrobot MK IV (FEI, Hillsboro OR) at 4 °C with 100% humidity, grids were blotted for 3-4 sec and plunged into liquid ethane. Grids were stored in liquid nitrogen prior to data collection.

CryoEM data was collected at the National Center for CryoEM Access and Training (NCCAT) on a Titan Krios microscope (ThermoFisher) operating at 300 kV equipped with energy filter (20 keV) and K3 detector (Gatan, Inc.). Micrographs were collected in a movie-mode with a total dose of 54 e^-^/Å^2^ spreading over 40 frames using Leginon at a nominal magnification of 81,000X in super-resolution mode, corresponding to a calibrated pixel size of 0.53 Å/pixel. The defocus values were set to range from -0.8 μm to -2.5 μm. Movie frames were aligned using Motioncor2 (44) and the contrast transfer function (CTF) parameters were estimated using GCTF (45). Particles were picked and 2D classified by cryoSPARC (46) and then exported for further processing in RELION (47). 3D-auto refinement was performed to generate the initial consensus map followed by mask-free 3D classification without alignment to further select good particles. High-quality particles were pools and used in alignment-free 3D classifications with a mask covering L1 stalk and TSV-IRES RNA. Particles from each class were exported to cryoSPARC for final reconstruction using the non-uniform refinement option.

### Model Building and Refinement

The coordinates were built from the available coordinate of the same ribosome (PDB ID: 7EWC) with approportionate adjustment and placement of hygromycin B in Chimera (48) or COOT (49). No water molecules were added. Manual examination and adjustment of the entire structure was carried out using COOT (49) before rounds of real-space refinement as implemented in PHENIX (50) were carried out until good map correlation coefficients and geometric values were reached (Supplementary Table 1).

### RADtool Analysis

We employed RADtool (Ribosome Analysis Database tool) (51) to compare conformations. For the reported structures, rotation angles were obtained using the RADtool as a plug-in to the VMD program and then were analyzed and compared against RAD database using the Web version of RADtool. The angles were extracted and plotted using Microsoft Excel.

## Acknowledgments

This work was supported by NIH grant R35 GM152081 to H.L. and NSF #2408763 to H.J.. Authors acknowledge the use of instruments at the Biological Science Imaging Resource supported by Florida State University and The Laboratory for BioMolecular Structure (LBMS). The Titan was funded from NIH grant S10 RR025080. The BioQuantum/K3 was funded from NIH grant U24 GM116788. The Vitrobot Mk IV was funded from NIH grant S10 RR024564. The Solaris Plasma Cleaner was funded from NIH grant S10 RR024564. The DE-64 was funded from NIH grant U24 GM116788. Some of this work was performed at the National Center for CryoEM Access and Training (NCCAT) and the Simons Electron Microscopy Center located at the New York Structural Biology Center, supported by the NIH Common Fund Transformative High Resolution Cryo-Electron Microscopy program (U24 GM129539,) and by grants from the Simons Foundation (SF349247) and NY State Assembly.

## Author Contributions

Y.Z., C.X., and H.L. designed the experiments. Y.Z. purified ribosomes, prepared cryo-EM grids, collected data and solved and modeled all structures, C.X. performed antibiotics sensitivity and thermostability analysis with the assistance of Y.Z.. X.C. and H.J. performed the cell-free ribosome activity assay, J. C. and H. J. designed and performed the cell-free translation assays, Y.Z., C.X., and H.L analyzed structure data and wrote the manuscript. All authors contributed to reviewing and editing the final manuscript.

## Data and materials availability

The atomic coordinates reported in this work are deposited to Protein Data Bank (PDB) with accession numbers 9DOV for 80S_hypoψ′_.IRES.HygB.I and ## for 80S_hypoψ′_.IRES.HygB.II. The cryoEM density map reported in this paper have been deposited to Electron Microscopy Data Bank (EMDB) with accession numbers EMD-47093 for 80S_hypoψ′_.IRES.HygB.I and EMD-### for 80S_hypoψ′_.IRES.HygB.II.

